# Dietary cystine restriction increases the proliferative capacity of the small intestine of mice

**DOI:** 10.1101/2023.08.10.552836

**Authors:** Judith C.W. de Jong, Kristel S. van Rooijen, Edwin C.A. Stigter, M. Can Gülersönmez, Marcel R. de Zoete, Janetta Top, Matthijs J.D. Baars, Yvonne Vercoulen, Folkert Kuipers, Saskia W.C. van Mil, Noortje Ijssennagger

## Abstract

Over 88 million people are currently estimated to have adopted towards a vegan or vegetarian diet. Cysteine is a semi-essential amino acid, which availability is largely dependent on dietary intake of meat, eggs and whole grains. Vegan/vegetarian diets are therefore inherently low in cysteine concentrations. Sufficient uptake of cysteine is crucial, as it serves as substrate for protein synthesis and conversion to taurine and glutathione. In this study, we therefore investigate the effect of low dietary cystine, the oxidized derivative of cysteine, on intestinal epithelial layer function. Mice (8/group) received a high fat diet with normal or low cystine concentration for 2 weeks. We observed no changes in plasma methionine, cysteine, taurine or glutathione levels after 2 weeks. Stem cell markers as well as the proliferation marker *Ki67* were increased upon cystine restriction in the small intestine. In line with this, gene set enrichment analysis indicated enrichment of Wnt signaling in the small intestine of mice on the low cystine diet, indicative of proliferative cells. Increased proliferation was absent in the colon. In the colon, dietary cystine restriction results in an increase in goblet cells, but no significant changes in the thickness of the mucus barrier or in its protective capacity. Also the microbiome was not changed upon dietary restriction. In conclusion, we show that cystine restriction for two weeks does not seem to induce any systemic effects. The increased proliferative capacity and number of goblet cells observed in the intestine may be the effect of starting epithelial damage or a reaction of the epithelium to start enlarging the absorptive capacity.

## Introduction

Cysteine is a semi-essential amino acid, which is provided by dietary intake of meat, eggs and whole grain. Cysteine can also be synthesized from the essential amino acid methionine via the transsulfuration pathway (1, 2). The majority of dietary cysteine is absorbed in the small intestine. Its ileal uptake occurs by the cystine/glutamate exchange transporter, xCT (3). Unabsorbed cysteine travels to the colon where it can be converted by sulfate- or sulfide-reducing bacteria to hydrogen sulfide. Cystine is the oxidized form of cysteine. Compared to cysteine, cystine from food sources is absorbed less efficient from the small intestine due to its lower digestibility (4). Intracellularly, cystine is reduced to cysteine by the NADH-dependent enzyme cystine reductase (5).

Next to protein synthesis, the three major fates of cysteine in the body are conversion to taurine and glutathione and hydrogen sulfide. These three metabolites have differential effects on intestinal function. Taurine is important for bile acid conjugation in the liver. Where in humans bile acids can be conjugated with either glycine or taurine (3:1), mice conjugate 95% of bile acids with taurine (6). Specific taurine-conjugated bile acids like taurocholic acid and taurolithocholic acid have been described to increase intestinal proliferation in intestinal cell lines (7, 8). Additionally, taurine-conjugated bile acids increase the abundance of sulfite-reducing bacteria B. Wadsworthia, which results in an increase in hydrogen sulfide production (9).

Low hydrogen sulfide concentration has been shown to be beneficial for integrity of the mucus layer (10), while excess hydrogen sulfide has been reported to cause breakdown of the mucus barrier and induce DNA damage in enterocytes (10, 11).

Lastly, glutathione (GSH) is an antioxidant which protects the intestine from oxidative stress and DNA damage. The addition of glutathione to intestinal porcine enterocytes on a cysteine-deprived medium restores proliferation and cell viability by replenishing the cysteine pool (12). Most of the above mentioned studies mimick high cystine/cysteine intake, with concurrent high concentrations of GSH, taurine and hydrogen sulfide, equivalent to a diet high in animal protein intake. As there is a trend towards adopting vegetarian and vegan diets, equivalent to low cystine availability (13), we questioned what the net effect of a low cystine diet is on intestinal function. We show that, in mice, cystine restriction results in an increase in proliferation and the number of stem cells associated with increased Wnt signaling, specifically in the small intestine. In addition, we observed that there is an increase in goblet cells and a trend towards an increase in mucus layer thickness in colon, however the protective capacity of the mucus layer is not changed.

## Results

### Dietary cystine restriction increases the expression of stem cell markers and proliferation specifically in the small intestine

Mice (n=8/group) received either a high fat (40 en%) diet or a high fat diet low in cystine (low cys) for two weeks. To investigate the effect of cystine restriction on the intestinal epithelial layer, gene expression levels of stem cell (*Lgr5, Sox9, Olfm4),* proliferative cell (*Ki67*), Paneth cell (*Lyz)*, enterocyte (*Krt20)* and goblet cell (*Muc2)* markers were determined in both small intestine (Fig 1A) and colon (Fig 1B) using qRT-PCR. Stem cell markers (*Lgr5, Sox9, Olfm4)* were or tended to be significantly increased upon cystine restriction. Markers of other cell types were not affected. The increase in stem cell markers was accompanied by higher Ki67 mRNA and protein expression, indicative of increased proliferation (Fig 1A, 1C). Moreover, the zone of Ki67+ cells was elongated as well upon cysteine restriction (Fig 1C). These effects on stem cell markers and proliferation were specific for the small intestine, as in the colon no changes were observed (Fig 1B).

**Figure 1:**
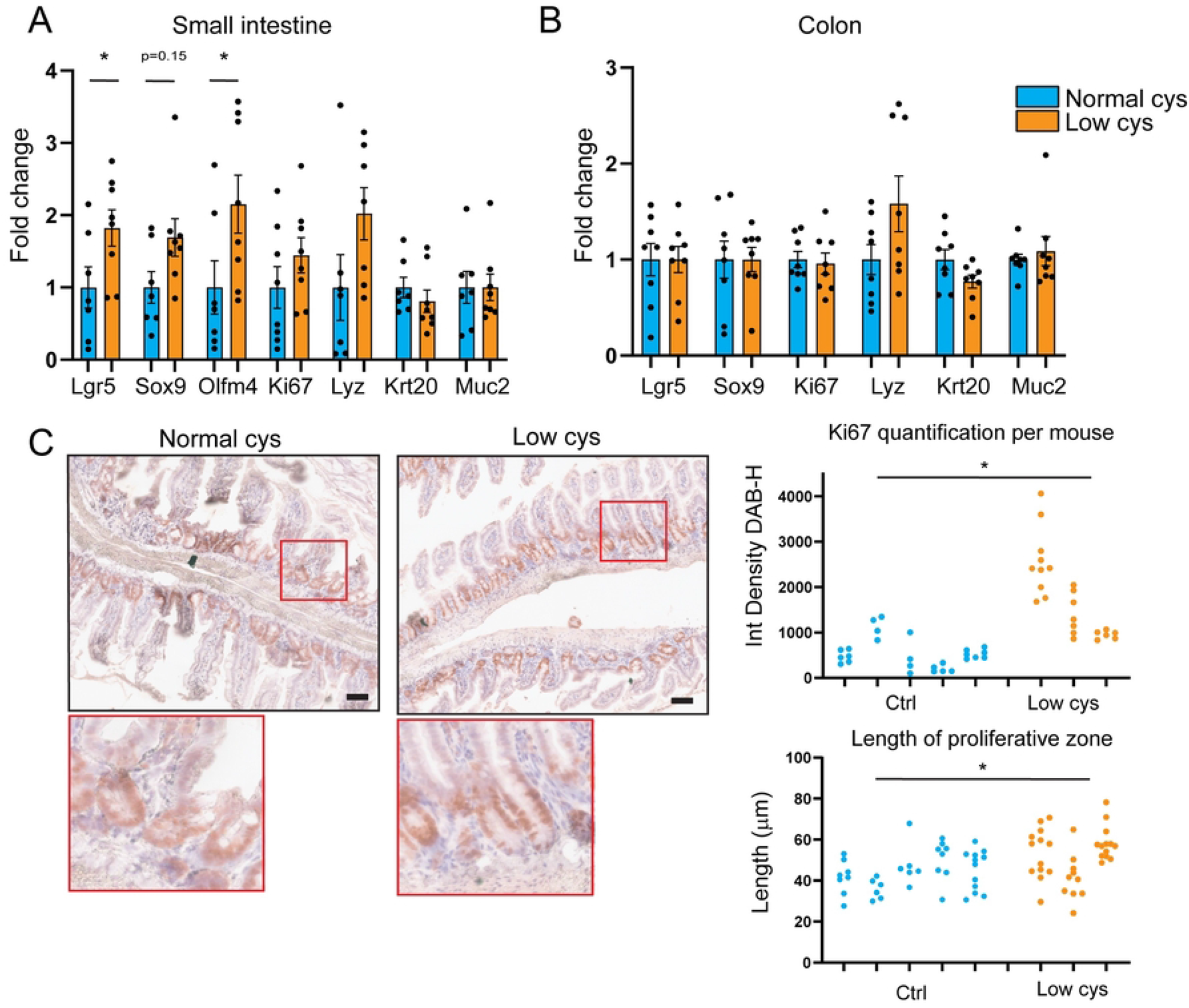
Dietary cystine restriction increases the expression of stem cell markers and proliferation specifically in small intestine A, B) qRT-PCR gene expression of markers of intestinal stem cell (Lgr5, Sox9, Olfm4), proliferative (Ki67), enterocyte (Krt20), Paneth cell (Lyz) and goblet cells (Muc2) normalized for normal cys (n=8/group, mean ± SEM, multiple Mann-Whitney tests, *p<0.05). C) (Left) Representative images of Ki67 immunohistochemistry on small intestinal crypts (scale bar = 50 µm). (Top Right) Quantification of integrated density of Ki67 corrected for haematoxylin of multiple crypts per mouse (unpaired t-test, *p<0.05). (Bottom Right) Quantification of the length of the proliferative zone stained by Ki67 of multiple crypts per mouse (unpaired t-test, *p<0.05).

### Dietary cystine restriction is not causing significant damage in the small intestine

An increase in proliferation and stem cells can be indicative of intestinal damage and the necessity to replenish the damaged cells (14, 15). However, since we did not observe any significant changes in body weight (Fig S1A), colon crypt length (Fig S1B) nor in expression of intestinal damage markers (*Ier3, Ripk3*, *Birc5,* Fig S1C & S1D) (16), a low cystine diet for 2 weeks did not seem to cause intestinal damage. However, we cannot exclude that the small increase in damage markers is a sign of beginning epithelial damage, which may become more severe upon longer cystine deprivation.

***Figure S1: Dietary cystine restriction is not causing significant damage in the intestine***

A) Difference in bodyweight of the mice between d14 and d0 (in gram) (n=8/group, mean ± SEM).

B) Total crypt length (µm) of 15 colonic crypts per mouse (with the median per mouse).

C, D) Gene expression of damage markers Ier3, Ripk3 and Birc5 in small intestine (C) and colon (D) (n=8/group, mean ± SEM).

### Dietary cystine restriction does not affect metabolism into cystine breakdown products

To unravel the underlying mechanism of the increased epithelial proliferation observed, we investigated the effects of cystine restriction on the metabolic pathways involving cysteine. The cystine-restricted diet did not cause a reduction in plasma cysteine levels (Fig 2A), suggesting that either the dietary deficiency is compensated for by methionine-to-cysteine conversion, or by a reduced production of taurine, glutathione and/or hydrogen sulfide. Plasma methionine levels were also not changed, neither was the mRNA expression of cystathionine γ-lyase (*CTH*), which is responsible for the last step in the conversion from methionine to cysteine (Fig 2A, 2B). This suggests that the methionine-to-cysteine conversion is not increased. The cysteine-restricted diet did not impact on glutathione production either, as both the plasma gluthathione concentration itself and mRNA expression of enzymes involved in the conversion from cystine to glutathione (*GSS, GCLC, GCLM)* were unchanged (Fig 2A, 2B). No other amino acids are affected either (table S2) suggesting that there is no cysteine deficiency after 2 weeks on a low cystine diet, or that the low cystine concentration (0.08%) still present in the low cys diet is sufficient.

**Figure 2:**
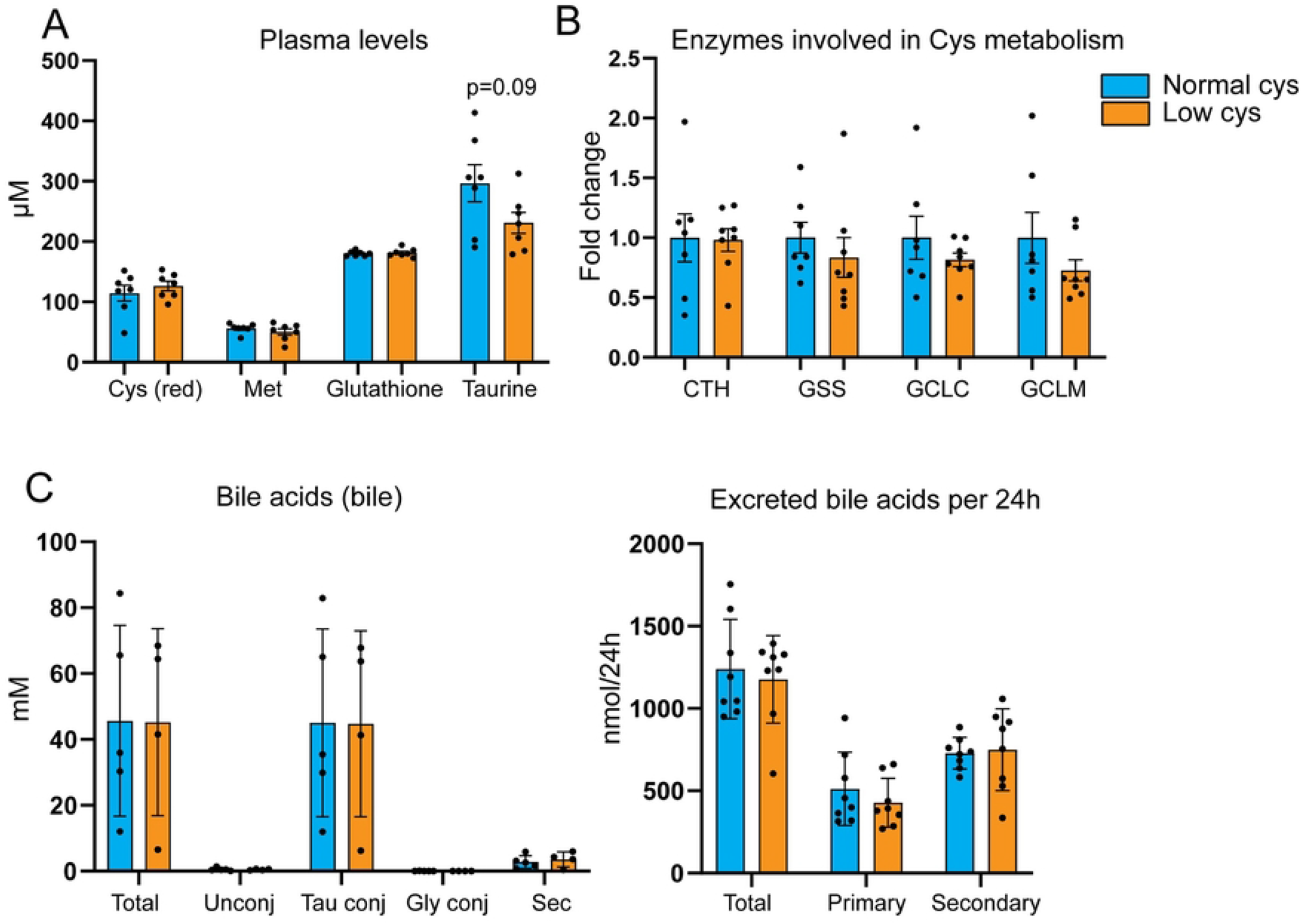
Dietary cystine restriction does not affect metabolism into cystine breakdown products A) Plasma cysteine (reduced), methionine, taurine and glutathione levels measured by metabolomics (n=8/group, mean ± SEM). B) qPCR gene expression of enzymes converting methionine to cysteine (CTH) and cysteine to glutathione (GSS, GCLC, GCLM) normalized by normal cys (n=8/group, mean ± SEM). C) Bile acid concentrations measured in bile (left) and excreted bile acids per 24h (right), measured in feces (n=8/group, mean ± SEM).

Cysteine restriction resulted in a trend towards decreased plasma taurine concentrations (Fig 2A). Methionine restriction has been shown to increase the ratio glycine to taurine conjugation of bile acids in mice (17). We therefore hypothesized that a low cystine diet could influence the bile acid conjugation ratio as well. Taurine and glycine conjugation of bile acids in gallbladder bile was not changed (Fig 2C), neither was total bile acid excretion into feces (Fig 2D). Taken together, we do not find evidence that cystine restriction affects the systemic availability of cysteine or its downstream catabolic metabolites.

**Table S2:** Concentrations of other amino acids measured using metabolomics.

|  | Normal cys (average) | Normal cys (SD) | Low cys (average) | Low cys (SD) |
| --- | --- | --- | --- | --- |
| Arginine (μM) | 88.5 | 16.8 | 90.7 | 11.2 |
| Glycine (μM) | 295.2 | 31.7 | 261.1 | 33.3 |
| Aspartic Acid (μM) | 12.6 | 2.0 | 12.6 | 2.4 |
| Citrulline (μM) | 56.7 | 5.8 | 55.1 | 10.0 |
| Glutamic Acid (μM) | 56.9 | 10.6 | 58.2 | 9.8 |
| Alanine (μM) | 422.7 | 83.6 | 376.3 | 120.6 |
| Tyrosine (μM) | 66.3 | 13.4 | 62.0 | 14.3 |
| Valine (μM) | 229.9 | 28.9 | 237.2 | 23.4 |
| Leucine (μM) | 152.0 | 21.1 | 153.9 | 23.3 |
| Phenylalanine (μM) | 61.3 | 7.8 | 59.1 | 6.6 |
| Histidine (μM) | 70.2 | 7.7 | 70.1 | 7.7 |
| Asparagine (μM) | 49.6 | 7.0 | 44.4 | 9.8 |
| Serine (μM) | 158.1 | 23.7 | 143.8 | 37.0 |
| Glutamine (μM) | 520.0 | 66.9 | 460.1 | 71.7 |
| Threonine (μM) | 90.3 | 6.4 | 88.8 | 9.1 |
| Proline (μM) | 91.1 | 13.1 | 84.2 | 17.3 |
| Ornithine (μM) | 46.6 | 18.6 | 46.3 | 5.3 |
| Lysine (μM) | 186.3 | 33.7 | 184.1 | 22.9 |
| Isoleucine (μM) | 100.3 | 17.2 | 98.1 | 6.9 |
| Tryptophan (μM) | 78.5 | 8.9 | 75.1 | 12.8 |

### Dietary cystine restriction increases Wnt signaling in small intestine

To further investigate the cause of increased proliferation in the small intestine upon cystine restriction, we performed RNA sequencing analysis on the small intestine. Dietary cysteine restriction leads to very little transcriptomic changes; only 16 genes were significantly differentially expressed when comparing low cysteine to normal cysteine (Fig S3). Most of these genes were upregulated in the low cysteine diet group, and represent either pseudogenes or genes part of the immunoglobulin kappa variable cluster (IGKV). However, gene set enrichment analysis (GSEA) revealed that one of the major pathways involved in intestinal stem cell homeostasis; Wnt signaling, was significantly enriched (FDR=0.07) upon a low cystine diet (Fig 3A). This was confirmed by qRT-PCR for the Wnt target genes *Axin2, Ascl2, Cmyc,* and *CyclinD1* (Fig 3B). As increased Wnt signaling is known to drive proliferation and is essential for stem cell maintenance, the increased Wnt signaling seen in cysteine-restricted mice, most likely causes the observed effects on stem cells and proliferation.

**Figure 3:**
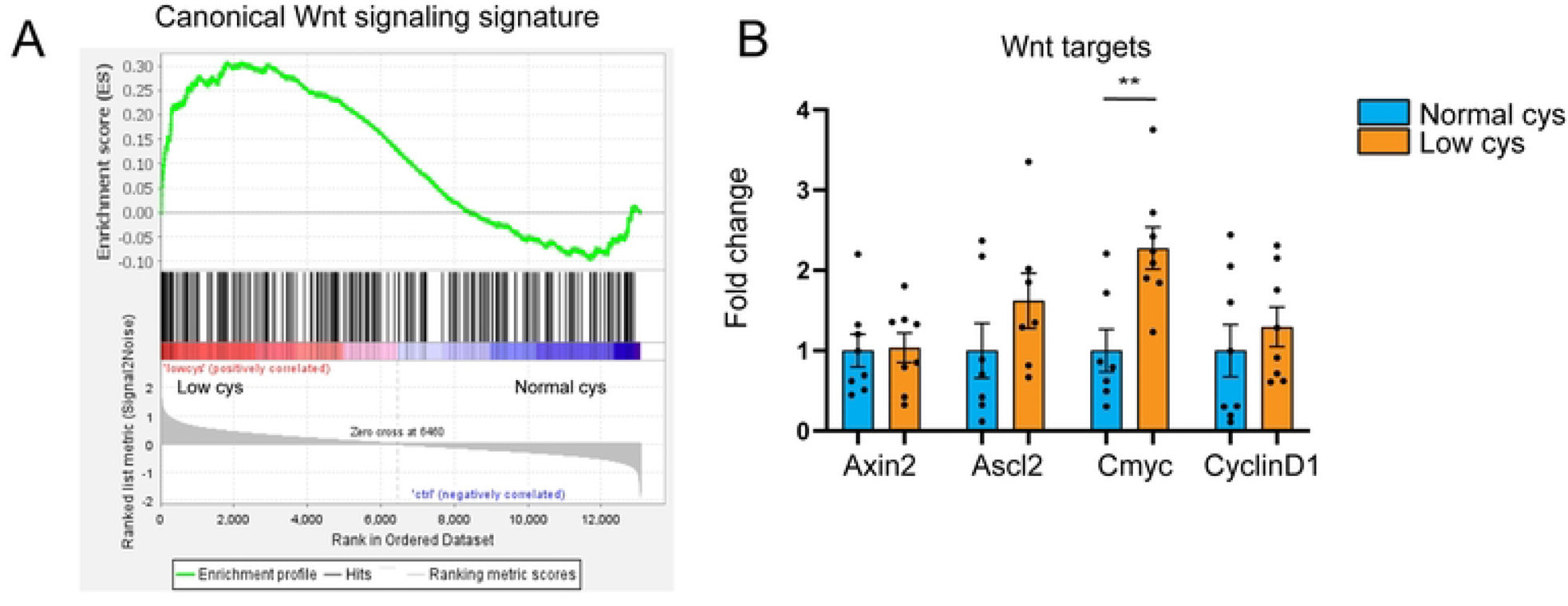
Dietary cystine restriction increases Wnt signaling in small intestine A) Gene set enrichment analysis for a Wnt signaling signature reveals significant enrichment with low cys vs ctrl (NES=1.37, FDR=0.07). B) qPCR gene expression analysis of Wnt target genes (n=8/group, mean ± SEM, multiple unpaired t-tests, **p<0.01).

***Figure S3: Significantly differentially expressed genes comparing low cys vs normal cys***

Heatmap of the 16 significantly differentially expressed genes comparing low cys to normal cys (fold change > 1.5, p-value < 0.05).

### Dietary cysteine restriction induces mucosal changes in the colon

The part of cystine that is not absorbed in the small intestine, eventually reaches the colon. The colonic epithelium is covered by mucus consisting of mucin polymers connected via disulfide bonds. This mucus layer limits the exposure of epithelial cells to toxins and bacteria. Certain bacteria such as sulfate reducing bacteria, can reduce these disulfide bonds (16), breaking the mucus barrier and increasing the exposure of epithelial cells to toxins and bacteria present in the lumen. Cystine also contains disulfide bonds which can be used as substrate for bacteria. Decreased luminal concentrations of cystine, such as in the low cys diet-treated mice, might therefore increase the chance that bacteria reduce the disulfide bonds from mucus rather than cystine. Therefore, we hypothesized that cystine restriction might affect the colonic mucus barrier. We observed that dietary cystine restriction resulted in an increase in the number of goblet cells and a trend towards an increase in the thickness of the secreted mucus layer (Fig 4A). Mucus barrier capacity, as measured by fluorescent bead penetration *ex vivo*, is not altered (Fig 4B). No differences in bacterial composition in colon were observed upon dietary restriction of cystine (Fig 4C), indicating that the effects on the mucus barrier were not an effect of changes in the abundance of mucus-degrading-or sulfate reducing bacteria.

**Figure 4:**
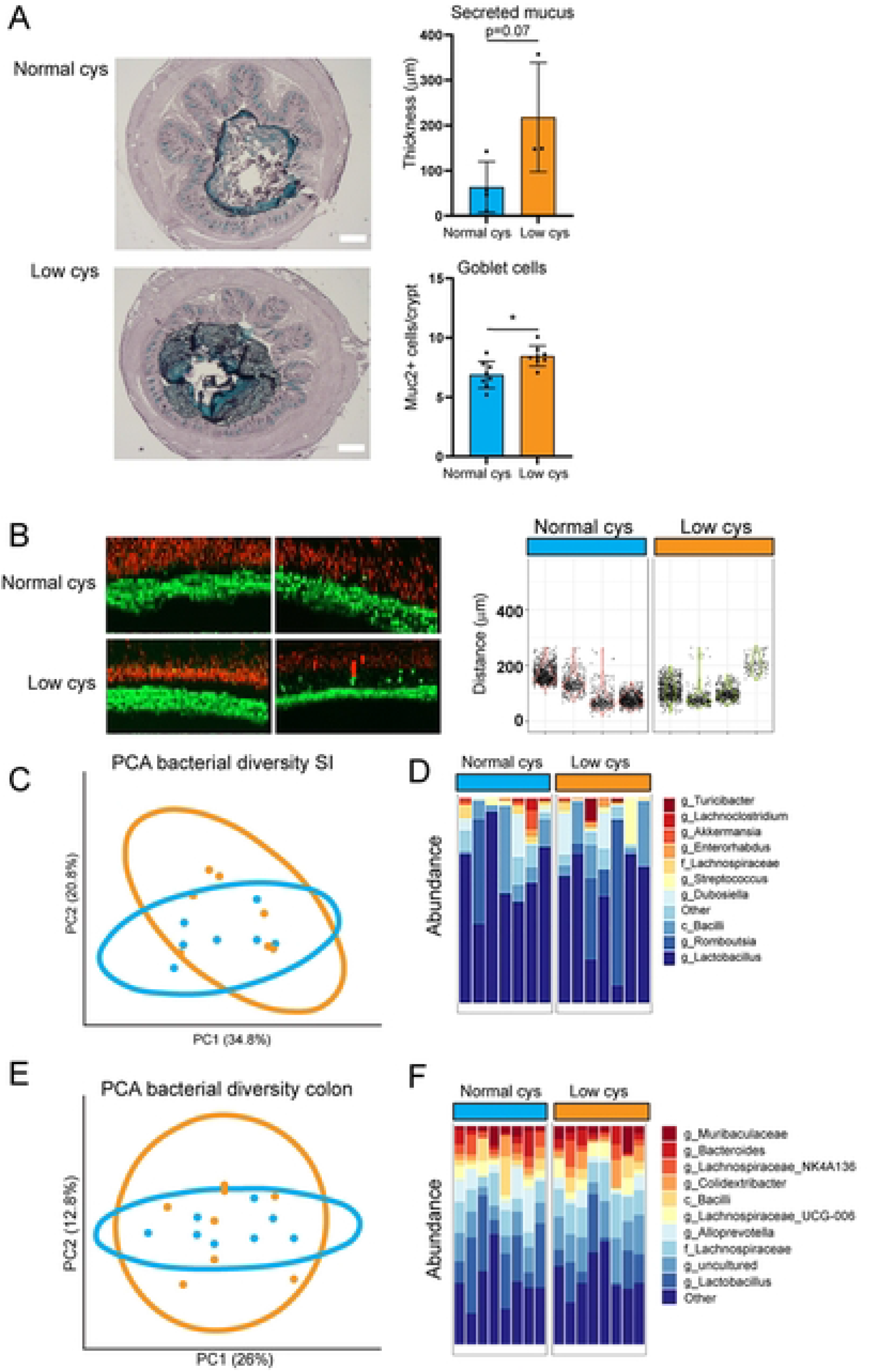
Dietary cystine restriction induces mucosal changes in the colon A) High Iron Diamine (HID) stainings for secreted mucus and quantifications (scalebar = 200 µm) and quantifications of Muc2+ cells per crypt (n=8/group, mean ± SEM, unpaired t-test, *p<0.05). B) Fluorescent bead microscopy for the mucus layer and quantifications of 4 mice per group (Each dot represents one bead) C) PCA plot of 16S sequencing of scrapings of small intestine. Each dot represents one mouse (n=7/group). D) Bacterial abundance analysis of the 16S sequencing for small intestine (n=7/group). E) PCA plot of 16S sequencing of scrapings of colon. Each dot represents one mouse (n=8/group). D) Bacterial abundance analysis of the 16S sequencing for colon (n=8/group).

## Discussion

Vegan and vegetarian diets contain limited amounts of cysteine, and although the human body can produce cysteine from methionine, the daily cysteine requirements are largely dependent upon direct dietary intake of cystine/cysteine. In this study, we therefore investigated the effect of dietary cystine restriction on the functionality of the intestinal cell types in mice. To our knowledge, this is the first study showing that cystine restriction affects intestinal proliferation and the expression of stem cell markers in the small intestine of mice, most probably via increased Wnt signaling. In addition, dietary cystine restriction led to an increase in goblet cells and a trend towards an increase in mucus layer thickness in the colon. Cystine restriction did not result in changes in plasma levels of cysteine, methionine, taurine and glutathione, indicating that there were no systemic effects of the low cystine diet. No changes in the microbiota were observed either, implying that the observed effects are most likely caused by the difference in cystine levels in the luminal content. The limiting systemic effects in our study could be attributed to several factors, including a relatively short duration of the diet (14 days) or the fact that the low cystine diet still contains 0.08% of cystine, which might not be limiting enough.

Certain bacteria such as sulfate reducing bacteria, reduce disulfide bonds present in colonic mucus (16), thereby breaking the mucus barrier and increasing the exposure of epithelial cells to toxins and bacteria present in the lumen. Cystine also contains disulfide bonds which specific bacteria use as substrate. Decreased luminal concentrations of cystine might therefore provide a shift in substrate usage by bacteria. This hypothesis is supported by a study reporting that cystine supplementation has a mucosal barrier enhancing effect (18). Whether this mechanism plays a role in this study needs to be further investigated, but the fact that there are more goblet cells in the colon in the low cystine group, even though the inner sterile layer is not thicker, might hint at a compensatory mechanism to produce more mucus.

Contradictory effects of cysteine, often in combination with methionine, on intestinal proliferation have been reported in diverse disease states and animal models. Both sulfur amino acid (methionine and cysteine) restriction and supplementation diets have been described to suppress intestinal proliferation in weaned or neonatal pigs after a 7-day intervention (19, 20). One study reports that supplementation with both methionine and cysteine decreases expression of β-catenin in the small intestine of weaned piglets, the downstream transcription factor in the Wnt pathway (19), after an intervention of 7 days, which is in line with our findings that Wnt signaling is increased with cystine restriction.

Additionally, multiple studies examine the effect of cystine in cancer models, since cystine has been reported to be essential for colorectal tumor growth. Cystine depletion has been described to reduce tumor xenograft growth in a mouse model (21) and metastases are sensitive to inhibition of cysteine uptake (22). Additionally, excessive production of hydrogen sulfide contributes to colorectal cancer development (23).

In conclusion, we show that cystine restriction for two weeks does not seem to induce any systemic effects. The increased proliferative capacity and number of goblet cells observed in the intestine may be the effect of starting epithelial damage or a reaction of the epithelium to start enlarging the absorptive capacity.

## Materials and methods

### Mice

The experiment was approved by the ethics committee of the University Medical Center Utrecht and was in accordance with European law. Eight-week-old male C57BL/6NRJ mice (Janvier) were housed individually in a room with controlled temperature (20–24°C), relative humidity (55% ± 15%), and a 12-h light–dark cycle. Mice were fed and had access to demineralized water ad libitum. Mice (n=8/group) received either the purified high fat (40 en%), low calcium AIN93-G diet (S9646-E070 Sniff, Germany) or the same diet in which the normally added 0.3% L-cystine was replaced equimolar by 0.11% alanine (S9646-E072). All mice were acclimatized for 1 week on the AIN93G HF control diet, before the 2-week intervention started.

Body weight was recorded during the intervention. Feces were quantitatively collected during the last 48 hours of the experiment and frozen at −20 °C for bile acid measurements. Mice were fasted 4 h before sacrifice. Periorbital puncture or heart puncture were performed to collect plasma after anesthesizing with isoflurane. Mice were sacrificed with cervical dislocation. Gall bladder including its content was collected and centrifuged at 10 000 RCF for 10 min to collect the bile, which was stored at −80 C.

The colon was excised, mesenteric fat was removed, and the colon was opened longitudinally, washed in PBS, and cut into 3 parts. The middle 1.5-cm colon tissue was formalin or carnoys fixed and paraffin embedded for histology. The remaining proximal and distal parts were scraped. Scrapings include the epithelial lining and lamina propria, but not the muscle layer. These scrapings were pooled per mouse, snap-frozen in liquid nitrogen, and stored at −80°C until further analysis. Colonic contents were sampled and snap-frozen for microbiota analysis. A similar procedure was followed for small intestine, which was divided in 4 parts equal in length. Of the last part of the small intestine (ileum), the first/proximal 1 cm was used for histology (swiss roll, formalin fixed), and the remaining part was scraped, snap-frozen in liquid nitrogen, and stored at −80°C until further analysis for mRNA expression.

### qRT-PCR

RNA was reverse transcribed using the iScript cDNA Synthesis Kit (Bio-Rad Laboratories BV, Veenendaal, The Netherlands). Real-time PCR was carried out using FastStart Universal SYBR Green Master Mix (Roche) on a CFX 384 Bio-Rad thermal cycler (Bio-Rad). mRNA expression of genes of interest were normalized to cyclophillin. Primer sequences can be found below (table 1).

**Table 1:**
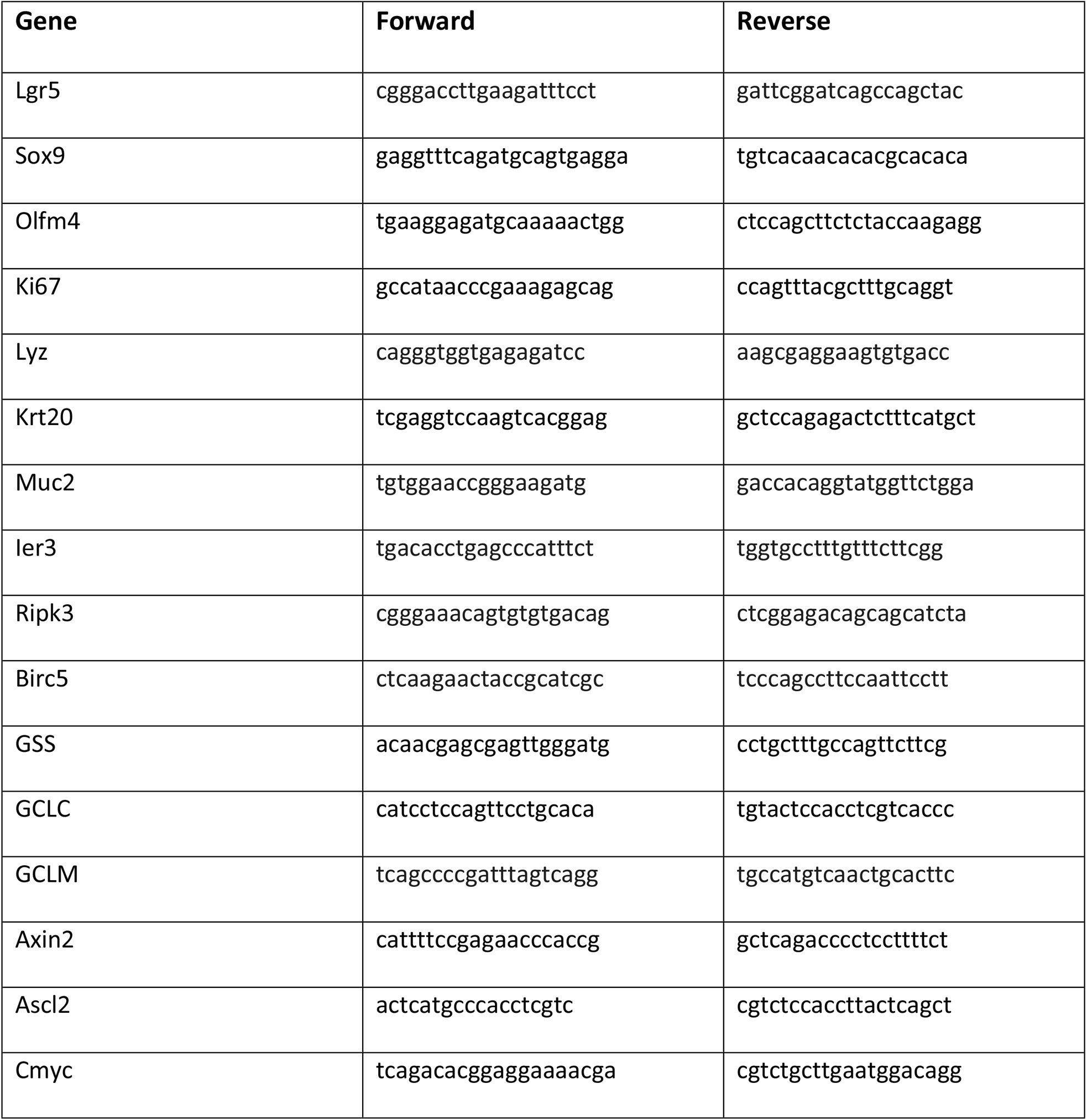
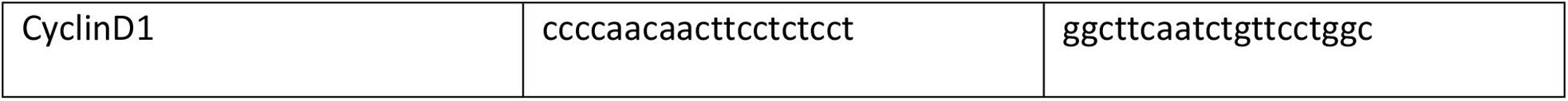
Primer sequences.

### Histology and immunohistochemistry

H&E staining was performed to assess the morphology of the tissue. To stain and quantify Ki67-positive cells, paraffin embedded colon sections (5 µm) were deparaffinized and rehydrated in a series of graded alcohols. Sections were incubated for 15 min in 3% H2O2 in PBS to block endogenous peroxidase activity. Sections were placed in antigen retrieval solution (sodium citrate buffer, pH=6) and heated in a microwave oven for 5 min 700W followed by 20 min 500W, after which they were cooled to room temperature. Sections were blocked with 10% normal goat serum (Sigma-Aldrich Chemie) in PBS-Tween 20 (0.05% v/v) for 30 min. Sections were then incubated with mouse anti-Ki67 (Dako, M724801-8, 1:200) or rabbit anti-Muc2 antibody (NBP1-31231, Novus Biologicals, USA, CO, 1:500) for 1h at room temperature. After washing, slides were incubated with the HRP-conjugated goat anti-mouse (1:200) for 30 min at room temperature. DAB substrate was used for HRP visualization and slides were counter stained with hematoxylin. Ki67 stainings quantification was done by deconvolution of the image in QuPath and measurement of mean intensity in ImageJ, normalized for the haematoxylin signal. 15 crypts per mouse were quantified. Crypt length was measured from the base of the epithelial layer until the bottom of the crypt using ImageJ.

For the High Iron Diamine staining (HID), staining sulfated mucins (brown) and carboxylated mucins (blue)), deparaffinized sections were incubated overnight in diamine solution. Then sections were incubated with alcian blue (pH-2.5) for 30 min. Sections were used to measure the thickness of the mucus layer (12 measurements for 3 mice per condition), measured from the surface top to the end of the mucus in the same orientation as the crypts.

### Plasma amino acids measurement

Organic solvents were ULC-MS grade and purchased from Biosolve (Valkenswaard, The Netherlands). Chemicals and standards were analytical grade and purchased from Sigma-Aldrich (Zwijndrecht, The Netherlands). Water was obtained fresh from a Milli Q instrument (Merck Millipore, Amsterdam, The Netherlands). A volume of 15 µL medium was transferred to a labelled 1.5 mL Eppendorf vial. A volume of 285 µL 80% acetonitrile also containing internal standards (final concentration 10 µM) was added and the sample was thoroughly mixed. The sample was centrifuged for 10 min at 17000xg in an Eppendorf centrifuge. A volume of 250 µL supernatant was transferred to a new, labelled Eppendorf vial and evaporated to dryness. Cell samples harvested in 300 µL ice-cold methanol were subjected to the same protocol. The residue was dissolved in 70 µL of borate buffer (pH 8.2) by thorough mixing and the derivatisation was started by adding 20 µL of AccQ-Tag reagent solution prepared according to the suppliers protocol (Waters, Etten-Leur, The Netherlands) after which the samples were vortex mixed and incubated at 55°C for 10 min. The sample was evaporated to dryness and the residue was dissolved in 120 µL 10% acetonitrile containing 0.1 mM formic acid and transferred to an LC sample vial. Analysis was performed on a system consisting of an Ultimate 3000 LC and an Thermo Scientific Q-Exactive FT mass spectrometer equipped with an HESI ion source (Thermo Scientific, Breda, The Netherlands). As a column a Waters HSS T3 (2.1×100 mm, 1.8 µm) was used, kept at a temperature of 40°C in the column oven. Eluent A used for analysis was milliQ water containing 0.1% formic acid, eluent B consisted of acetonitrile containing 0.1% formic acid. The LC gradient used for separation commenced by injecting 1 µL of sample and started at 0% B for 5 min. followed by a 10 min linear gradient to 75% B. In 0.5 min the gradient increase to 100% B and kept there for 1 min before returning to 0% B. The column was allowed to regenerate for 2.5 min prior to a next analysis. Total runtime was 18 min; flow rate was 400 µL/min. The mass spectrometer was operated in ESI-positive mode, full scan 100 – 1000 m/z, capillary temperature 300°C, sheath gas: 35, aux gas:2, resolution 30000, capillary voltage 3 kV.

### Bile acid measurements

Gall bladder BAs were measured as described in (24). Fecal BAs were measured as described in (25). In short, bile samples were diluted with ammonium acetate buffer 15 mM (pH = 8.0): (acetonitrile/methanol=75/25 v/v) = 50:50, v/v. A mixture of internal standards in methanol was added to the samples to reach a concentration of 2.5 µM. Both fecal and gall bladder BAs were quantified using an Ultra Performance Liquid Chromatography-Mass Spectrometry system (UPLC-MS2, Acquity H-Class Bio UPLC from Waters).

### RNA isolation and sequencing

Total RNA from colon and SI was isolated for qPCR and sequencing, using TRIzol reagent (Invitrogen) according to the manufacturer’s protocol. The RNA was further purified using either RNeasy Minikit columns (Qiagen) or the Nucleospin RNA mini kit (Macherey Nagel). For sequencing, libraries were prepared using Truseq RNA stranded polyA (Illumina) and sequenced on an Illumina Novaseq6000 in paired-end 50 bp reads. Quality control on the sequence reads from the raw FASTQ files was done with FastQC (v0.11.8). TrimGalore (v0.6.5) as used to trim reads based on quality and adapter presence after which FastQC was again used to check the resulting quality. rRNA reads were filtered out using SortMeRNA (v4.3.3) after which the resulting reads were aligned to the reference genome fasta (Mm_GRCm38_gatk_sorted.fasta) using the STAR (v2.7.3a) aligner. Followup QC on the mapped (bam) files was done using Sambamba (v0.7.0), RSeQC (v3.0.1) and PreSeq (v2.0.3). Readcounts were then generated using the Subread FeatureCounts module (v2.0.0) with the Mus_musculus.GRCm38.70.gtf gtf file as annotation, after which normalization was done using the R-package edgeR (v3.28). Differential Expression analysis was performed with an inhouse R-script using DESeq2 (v1.28) taking the raw readcounts as input. Finally a summary report was created using MultiQC (v1.9). Gene set enrichment analysis (GSEA) was performed using the GSEA tool from the Molecular Signature Database (MSigDB), a joint project of UC San Diego and the Broad Institute ((26, 27). Sequencing data has been made available in the NCBI Gene Expression Omnibus (GEO) under accession number GSE234566.

### Fluorescent beads penetration assay

Mucus barrier function was determined in C57BL/6NRJ mice (Janvier) on the low cys (n=4) or the normal diet (n=4) (same diets and conditions as described above). Mice were sacrificed and a 1 cm piece of colonic tissue, 2 cm above the rectum, was dissected for mucus measurements. This assay was performed as described in Ijssennagger et al. (2021) (24). In short, tissue was visualized using a Syto9 green fluorescent nucleic acid stain (ThermoFisher) at the apical side. After 10 mins, FluoSphere Crimson microbeads (1 µm, ThermoFisher) diluted 30 times in Krebs mannitol buffer were added on top of the colon tissue, to measure mucus penetrability. The tissue and beads were visualized using a Zeiss AxioImager Z1 (Carl Zeiss, Germany). Quantifications were done using Fiji and represent the distance of the beads to the cell monolayer.

### Bacterial DNA extraction and 16S sequencing

DNA extraction was performed using a modified protocol of the QIAamp fast DNA stool mini kit (Qiagen, Venlo, the Netherlands) as previously described (28, 29). In brief, 0.2 g feces was added to ‘lysing matrix A, 2 ml tubes’ (MP biomedicals, Landsmeer, the Netherlands) containing 1 ml InhibitEx buffer (Qiagen). Two rounds of bead incubations were applied at 3.5 m/s for 2 min, followed by 2 min incubation on ice using the FastPrep-24 (MP biomedicals). After 7 min of incubation at 95 °C, the protocol of the fast DNA stool mini kit protocol (Qiagen) was followed from the proteinase K treatment step onwards. Total DNA was quantified by Picogreen assay (Thermo Fisher Scientific, Waltham, MA, USA). The 469 bp V3 and V4 hyper-variable regions of the 16S rRNA gene were amplified and sequenced using the Illumina MiSeq instrument and Reagent Kit v3 (600-cycle) according to Fadrosh et al. (30). Negative controls and mock communities (ZymoBIOMICS microbial community standard (D6300) and ZymoBIOMICS microbial community DNA standard (D6305), ZymoBIOMICS Microbial Community Standard II (Log Distribution) (D6310), Zymo research, USA) were used from the beginning of DNA isolation up to the data analysis stage and matched with the distribution expected mock compositions. For analysis, the QIIME2 microbial community analysis pipeline (version 2021.4) (31) was used with DADA2 for sequence variant detection (with default settings, except for --p-trunc-len-f 255 --p-trunc-len-r 240) (32), and SILVA as 16S rRNA reference gene database (SILVA 138) (33). Sequencing data has been made available on the European Nucleotide Archive under project PRJEB63227.

## Acknowledgements

We thank the Utrecht Sequencing Facility (USEQ, UMC Utrecht) for performing RNA sequencing; F. Mulder (UMC Utrecht) for doing the RNAseq analyses; M. Koehorst for bile acid analysis; L. Kleij for microscopy assistance and R. van der Meer for scientific discussions.

## Financial Support

This work was financially supported by DSM Nutritional Products and the Dutch Technology Foundation STW (grant number 14940), which is the Applied Science Division of NWO, and Technology Programme of the Ministry of Economic Affairs. NI is further financially supported by the MLDS Career Development Grant (CDG16-04) and by the Wilhelmina Children’s Hospital Research Fund. SvM is further financially supported by the Netherlands Organisation for Health Research and Development (ZonMW; VICI grant, no: 09150181910029 and Aspasia grant, no: 015.015.013).

## Author contributions

Conceptualization: Noortje Ijssennagger, Saskia W.C. van Mil; methodology and investigation: Judith W. de Jong, Kristel van Rooijen, Noortje Ijssennagger, Saskia W.C. van Mil; metabolomics: Edwin C.A. Stigter, M. Can Gülersönmez; bacterial analyses: Janetta Top, Marcel R. de Zoete; mucus barrier visualization: Noortje Ijssennagger, Kristel van Rooijen, Matthijs J.D. Baars, Yvonne Vercoulen; Bile acids measurements; Noortje Ijssennagger, Folkert Kuipers; manuscript writing: Judith C.W. de Jong, Noortje Ijssennagger, Saskia W.C. van Mil; review and editing: Judith C.W. de Jong, Noortje Ijssennagger, Saskia W.C. van Mil; funding acquisition: Noortje Ijssennagger, Saskia W.C. van Mil.

## Declaration of competing interest

The authors declare that they have no known competing financial interests or personal relationships that could have appeared to influence the work reported in this paper.

## Notes

### Competing Interest Statement

The authors have declared no competing interest.

